# Antiviral activity of influenza A virus defective interfering particles against SARS-CoV-2 replication *in vitro* through stimulation of innate immunity

**DOI:** 10.1101/2021.02.19.431972

**Authors:** U. Rand, S.Y. Kupke, H. Shkarlet, M.D. Hein, T. Hirsch, P. Marichal-Gallardo, L. Cicin-Sain, U. Reichl, D. Bruder

## Abstract

SARS-CoV-2 causing COVID-19 emerged in late 2019 and resulted in a devastating pandemic. Although the first approved vaccines were already administered by the end of 2020, worldwide vaccine availability is still limited. Moreover, immune escape variants of the virus are emerging against which the current vaccines may confer only limited protection. Further, existing antivirals and treatment options against COVID-19 only show limited efficacy. Influenza A virus (IAV) defective interfering particles (DIPs) were previously proposed not only for antiviral treatment of the influenza disease but also for pan-specific treatment of interferon (IFN)-sensitive respiratory virus infections. To investigate the applicability of IAV DIPs as an antiviral for the treatment of COVID-19, we conducted *in vitro* co-infection experiments with cell culture-derived DIPs and the IFN-sensitive SARS-CoV-2 in human lung cells. We show that treatment with IAV DIPs leads to complete abrogation of SARS-CoV-2 replication. Moreover, this inhibitory effect was dependent on janus kinase/signal transducers and activators of transcription (JAK/STAT) signaling. Further, our results suggest boosting of IFN-induced antiviral activity by IAV DIPs as a major contributor in suppressing SARS-CoV-2 replication. Thus, we propose IAV DIPs as an effective antiviral agent for treatment of COVID-19, and potentially also for suppressing the replication of new variants of SARS-CoV-2.

## Introduction

Severe acute respiratory syndrome coronavirus 2 (SARS-CoV-2), causing coronavirus disease 2019 (COVID-19), poses a severe burden to public health, economy and society. To date, almost four million deaths were reported in the context of SARS-CoV-2 infection (WHO, covid19.who.int). Starting early 2020, there has been a unprecedented race for the development of novel vaccines, their production, and execution of safety and immunogenicity studies in clinical trials (Krammer, 2020, Walsh et al., 2020, Voysey et al., 2020, Jackson et al., 2020, Logunov et al., 2020, Palacios et al., 2020). These efforts led to the vaccination of the first individuals outside clinical trials in late 2020. While vaccination typically provides the best protection against virus infections and disease onset, the worldwide manufacturing capacity of COVID-19 vaccines, and the infrastructure required for vaccination are still limited. In addition, vaccination is a prophylactic measure and not applicable for therapeutic treatment of acute infections. Therefore, as an alternative option, the development of antivirals for treatment of COVID-19 is essential. Yet, remdesivir (in clinical use) showed only limited efficacy (Wang et al., 2020, Beigel et al., 2020, Pan et al., 2020), while other repurposed drug candidates (e.g., hydroxychloroquine and lopinavir-ritonavir) showed a lack of efficacy (Boulware et al., 2020, Cao et al., 2020). In addition, corticosteroids (i.e., dexamethasone (Tomazini et al., 2020)) and cocktails of monoclonal antibodies (e.g., bamlanivimab (Chen et al., 2020b)) are used in the clinic and show an antiviral effect. However, the appearance of new SARS-CoV-2 variants poses a constant risk to lose efficacy of highly specific treatments, including vaccination and therapeutic antibodies. Thus, there is a need to develop more broadly acting, cost-effective antivirals that ideally are easily scalable in production.

Influenza A virus (IAV) defective interfering particles (DIPs) were previously proposed for antiviral treatment against IAV infections (Zhao et al., 2018, Vignuzzi and Lopez, 2019, Meir et al., 2020, Tapia et al., 2019, Yang et al., 2019, Harding et al., 2019, Tanner et al., 2016, Huo et al., 2020b, Kupke et al., 2020), but also for pan-specific treatment of other respiratory viral diseases (Dimmock and Easton, 2014, Dimmock and Easton, 2015). IAV DIPs typically carry a large internal deletion in their genome, rendering them defective in virus replication (Alnaji and Brooke, 2020, Ziegler and Botten, 2020, Andreu-Moreno and Sanjuan, 2020). Furthermore, DIPs suppress and interfere specifically with homologous viral replication in a co-infection scenario, a process known as replication interference. As a result, administration of IAV DIPs to mice resulted in full protection against an otherwise lethal IAV infection (Dimmock et al., 2008, Huo et al., 2020b, Hein et al., 2021c, Hein et al., 2021a). In the ferret model, treatment of IAV-infected animals resulted in a reduced severity of disease pathogenesis (Dimmock et al., 2012). Intriguingly, mice were also protected against a lethal infection with the unrelated influenza B virus (Scott et al., 2011) and pneumonia virus of mice (PVM) from the family *Paramyxoviridae* (Easton et al., 2011). Here, protection was not attributed to replication interference but to the ability of IAV DIPs to stimulate innate immunity.

SARS-CoV-2 replication seems to modulate and inhibit the interferon (IFN) response in infected target cells (Chen et al., 2020a, Konno et al., 2020, Lei et al., 2020). Still, it was also shown to be susceptible to inhibition by exogenously added IFNs *in vitro* (Busnadiego et al., 2020, Felgenhauer et al., 2020), *in vivo* (Hoagland et al., 2021) and in clinical trials (Monk et al., 2020). Therapies using recombinant IFNs, however, are cost intensive and pose the risk of unwanted side effects including the formation of auto-antibodies against the cytokine (reviewed in (Sleijfer et al., 2005)). To prevent this, we wondered whether IAV DIPs would be able to suppress SARS-CoV-2 replication through their ability to stimulate a physiological IFN response in target cells. To test this, we produced two promising candidate DIPs, a prototypic, well-characterized conventional IAV DIP “DI244” (Dimmock and Easton, 2014), and a novel type of IAV DIP “OP7”, that contains point mutations instead of a large internal deletion in the genome (Kupke et al., 2019), using a cell culture-based production process (Hein et al., 2021c, Hein et al., 2021a, Hein et al., 2021b).

Here, we used Calu-3 cells (human lung cancer cell line) for *in vitro* co-infection experiments with SARS-CoV-2 and DI244 or OP7, respectively. Both DIPs were able to completely inhibit SARS-CoV-2 replication and spreading in a range comparable to IFN-β or remdesivir treatment. Moreover, we show that the inhibitory effect of IAV DIPs was due to their ability to induce innate immune responses that signal via janus kinase/signal transducers and activators of transcription (JAK/STAT). In addition, our results show that IAV DIP infection triggers elevated host cell type-I and type-III IFN production and subsequent IFN-induced antiviral activity. Thus, we propose IAV DIPs as effective antiviral agents for the treatment of COVID-19 and, potentially as universal antiviral agents not only against different influenza subtypes but also against other (including newly emerging) IFN-sensitive respiratory viruses.

## Results

### SARS-CoV-2 replication is abrogated by IAV DIP treatment *in vitro*

In order to test the antiviral efficacy of IAV DIPs on replication of SARS-CoV-2, we conducted *in vitro* co-infection experiments in Calu-3 cells. For this, we infected cells with SARS-CoV-2 (multiplicity of infection (MOI) = 0.03) and highly concentrated IAV DIPs (DI244 or OP7), respectively, from cell culture-based production and membrane-based chromatographic purification (Marichal-Gallardo et al., 2017, Hein et al., 2021c, Hein et al., 2021a). At 3 days post infection (dpi), cells were stained for the SARS-CoV-2 spike (S) protein (Fig. 1A). Indeed, S protein expression was significantly reduced in cells co-treated with DI244 or OP7 compared to cells infected with SARS-CoV-2 only, indicating suppression of SARS-CoV-2 replication by IAV DIP co-infection. In agreement with this observation, images from live-cell microscopy show a clearly reduced cytopathic effect upon DIP co-infection from ~36 hpi on (Fig. 1B).

**Fig. 1.**
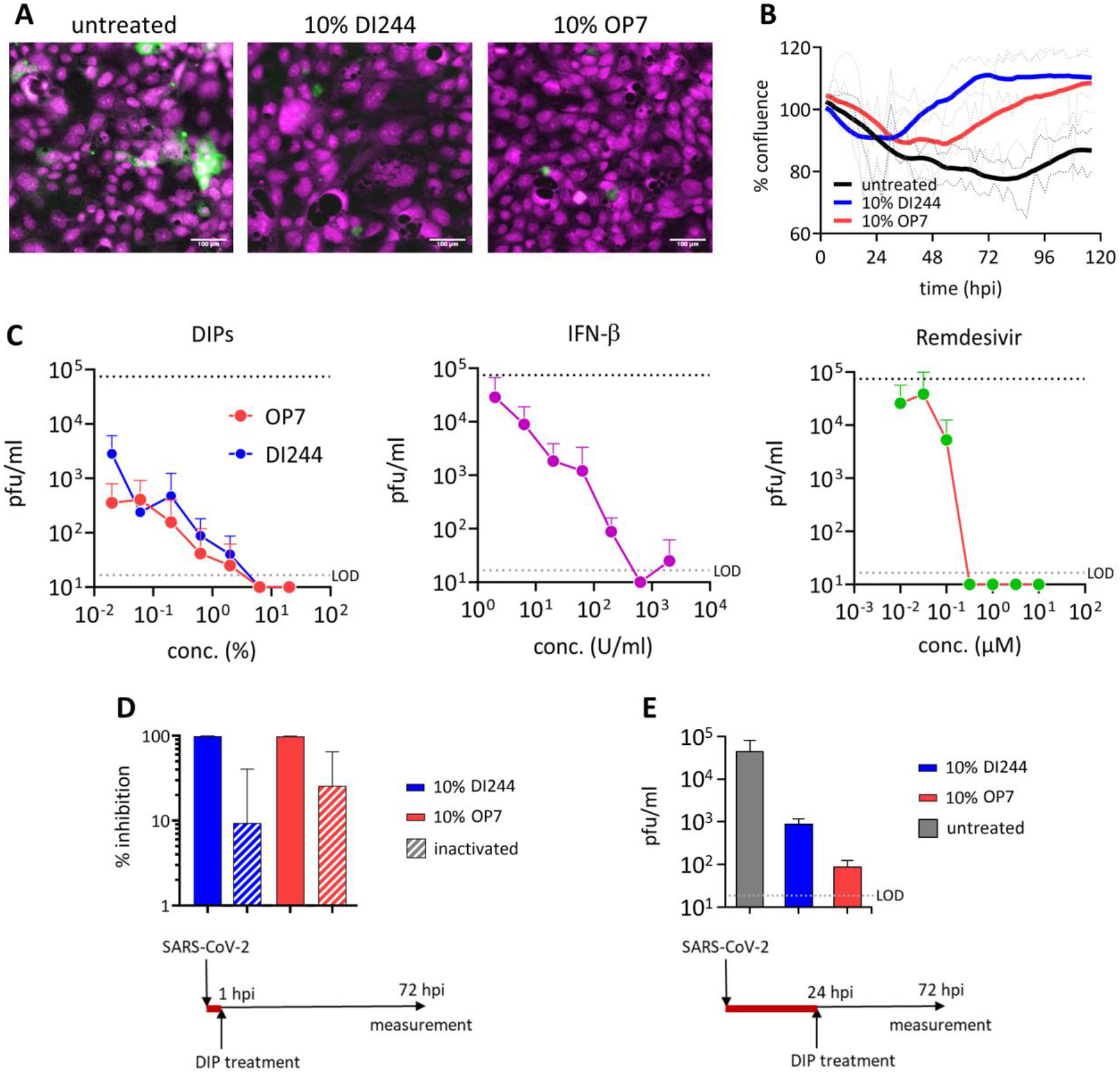
Inhibition of SARS-CoV-2 replication and spreading by IAV DIPs. SARS-CoV-2-infected Calu-3 cells (MOI=0.03) were treated with IAV DIPs (DI244 or OP7), IFN-β, or remdesivir at 1 hour post infection (hpi). For DI244 and OP7 treatment, chromatographically purified and highly concentrated cell culture-derived DIP material (Hein et al., 2021c, Hein et al., 2021a) was used. % (v/v) indicates the fraction with respect to the cell culture volume of 100 μL. Stock concentration, 5.6 × 10^8^ and 1.12 × 10^11^ DI vRNAs/mL for DI244 and OP7, respectively. (**A**) Immunofluorescence analysis of the SARS-CoV-2 S protein expression (green, magenta: DNA) at 3 dpi. Scale bar, 100 μm. (**B**) Cytopathic effect. Confluence (% of initial) was measured by live-cell microscopy at 2 h intervals. Thick lines represent smoothened data (Savitzky-Golay filter), dotted lines show SD of original data (n=2, independent experiments). (**C**) Effective concentration range of DI244 and OP7 compared to IFN-β and remdesivir. Viral titers were determined from the supernatant at 3 dpi by plaque assay. Upper dotted line indicates virus titer in untreated cells, lower dotted line shows the LOD. Independent experiments were conducted; mean ± SD (n=3) is shown. (**D**) SARS-CoV-2 growth inhibition by active and inactive DIPs. SARS-CoV-2 infected cells were treated with active or UV-inactivated DIPs at 1 hpi. Percentage inhibition of viral growth relative to mock treatment is shown; mean ± SEM (n=4) is depicted. (**E**) DIP superinfection 24 h post SARS-CoV-2 infection. Independent experiments were conducted; mean ± SD (n=2) is shown.

In order to test the antiviral efficiency of IAV DIPs in comparison with clinically relevant antivirals, dose-effect curves were generated by testing different concentrations of IAV DIPs for the treatment of SARS-CoV-2-infected cells (Fig. 1C). As a read-out for SARS-CoV-2 replication, we determined the infectious titer from supernatants at 3 dpi using plaque assays with Vero-6 cells. Please note that infection with only defective, replication-incompetent IAV DIPs does not result in the release of progeny virions, as demonstrated by negative plaque titers (Hein et al., 2021c, Hein et al., 2021a). Strikingly, SARS-CoV-2 replication was severely diminished upon IAV DIP co-infection. In particular, at high DI244 and OP7 concentrations, no SARS-CoV-2 plaque titers were detectable anymore, while untreated cells showed a titer of 7.5 x 10^4^ plaque-forming units (pfu)/mL. Suppression of SARS-CoV-2 replication decreased with increasing dilution of DIPs. Remarkably, though, the treatment with both DIPs at a dilution of 1:1000 still resulted in a pronounced inhibition of SARS-CoV-2 replication, corresponding to a 26-fold and 210-fold reduction in plaque titers for DI244 and OP7 treatment, respectively. For comparison, we also tested the inhibitory capacity of IFN-β or remdesivir treatment on SARS-CoV-2 replication in infected target cells. Both agents were also able to diminish SARS-CoV-2 plaque titers to below the limit of detection (LOD), until a concentration of 633 U/mL for IFN-β and 0.32 μM for remdesivir. Yet, inhibiting effects ceased significantly faster with increasing dilutions, most apparently observed for remdesivir, for which treatment with a concentration of 0.03 μM (1:10 dilution) already did not result an inhibitory effect anymore.

Fig. 1D illustrates residual SARS-CoV-2 inhibition caused by inactivated DIPs. These DIPs were previously UV-irradiated until no interfering efficacy against IAV replication was observed anymore *in vitro* (Hein et al., 2021c, Hein et al., 2021a), indicating the degradation of the causative interfering agent, i.e., the genomic defective interfering (DI) viral RNA (vRNA). The finding that inhibition caused by active DIPs was more efficient suggests a specific interfering activity of active IAV DIPs with SARS-CoV-2 replication. Of note, active DIPs still conferred a pronounced antiviral effect even when applied 24 h after preceding SARS-CoV-2 infection (Fig. 1E).

In conclusion, treatment with both DI244 and OP7 IAV DIPs completely abolished SARS-CoV-2 replication during *in vitro* co-infections. While the inhibitory potential was comparable to IFN-β and remdesivir treatment, the antiviral effects of IAV DIPs were more sustained with increasing dilution.

### IAV DIP infection enhances type-I and type-III IFN responses and inhibit SARS-CoV-2 replication via Janus-kinase signaling

Next, to investigate our hypothesis whether inhibition of SARS-CoV-2 replication by DIPs was due to their ability to stimulate the IFN system, we used ruxolitinib in co-infection experiments. This small molecule drug is an efficient inhibitor of JAK, which are key effectors in the IFN system. Upon IFN sensing, JAKs typically recruit STATs, ultimately leading to the upregulation of IFN-stimulated gene (ISGs). ISGs encode for effector molecules that limit viral replication by inducing an antiviral state in the infected as well as uninfected neighboring cells. Fig. 2 shows the results of SARS-CoV-2 and IAV DIP co-infection upon treatment with ruxolitinib. While DI244 and OP7 co-infection almost completely inhibited SARS-CoV-2 replication, additional treatment with ruxolitinib abrogated the suppressive effect of both IAV DIPs. Specifically, virus titers under JAK signaling inhibition were comparable to SARS-CoV-2 infection in the absence of DIPs. These results suggest a major contribution of unspecific innate immune activation by IAV DIPs in interfering with SARS-CoV-2 replication.

**Fig. 2.**
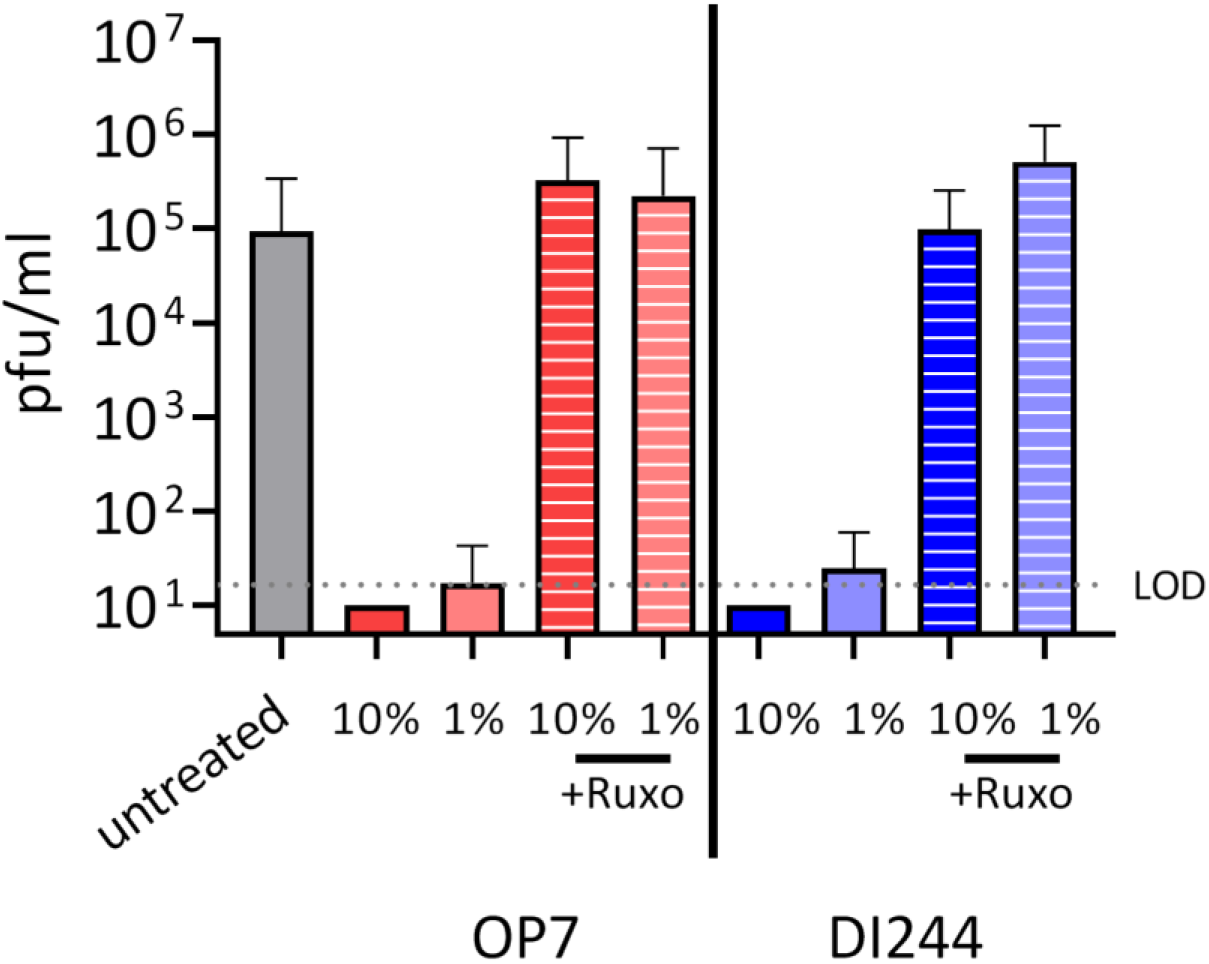
Suppression of SARS-CoV-2 replication by IAV DIPs under JAK inhibition. SARS-CoV-2-infected Calu-3 cells (MOI=0.03) were co-infected with IAV DIPs (DI244 or OP7) at 1 hpi in the presence of absence of ruxolitinib (JAK inhibitor). % (v/v) indicates the fraction of DIPs (highly concentrated cell culture-derived material (Hein et al., 2021c, Hein et al., 2021a) with respect to the cell culture volume of 100 μL. Viral titers were determined from the supernatant at 3 dpi by plaque assay. Dotted line shows the LOD. Independent experiments were conducted; mean ± SD (n=4) is depicted.

In order to shed light onto correlates of IAV DIP-mediated SARS-CoV-2 suppression, we used real-time RT-qPCR for quantification of gene expression to assess innate immune responses during co-infections in more detail (Fig. 3A). We observed a significant upregulation of type I and III IFN (i.e., *IFNB1* and *IFNL1*, respectively) expression at early times following IAV DIP and SARS-CoV-2 co-infection compared to SARS-CoV-2 single infection. Specifically, we detected one to two log_10_ higher *IFNB1* and *IFNL1* mRNA levels at 6 hpi. In addition, canonical antiviral gene expression, indicated by *MX1* and *RSAD2,* was elevated. This early upregulation of innate immune responses may well explain the strong antiviral activity of DI244 and OP7 against SARS-CoV-2 replication. In agreement with this, we detected significant uptake of the non-replicating DI244 and OP7 vRNA (~10E3 molecules/cell) (Fig. 4A) and concomitant upregulation of *DDX58* (Fig. 4B), encoding for retinoic acid inducible gene I (RIG-I); a pattern recognition receptor (PRR) critical for detection of viral nucleic acids and initiation of cellular antiviral responses. The significantly lower levels of DI vRNAs detected for cells treated with inactive DIPs can be explained by their efficient degradation by the inactivation procedure (i.e., UV-irradiation). As a control, we infected cells with only active or inactive IAV DIPs. Here, only active DIPs were able to mount an elevated innate immune response (Fig. 3A, lower panels). Interestingly, canonical antiviral gene expression (*MX1* and *RSAD2*) was enhanced until 72 hpi for the active DIP-only treatment, implying that DI244 and OP7 can potentially raise a long-lived antiviral ISG response.

**Fig. 3.**
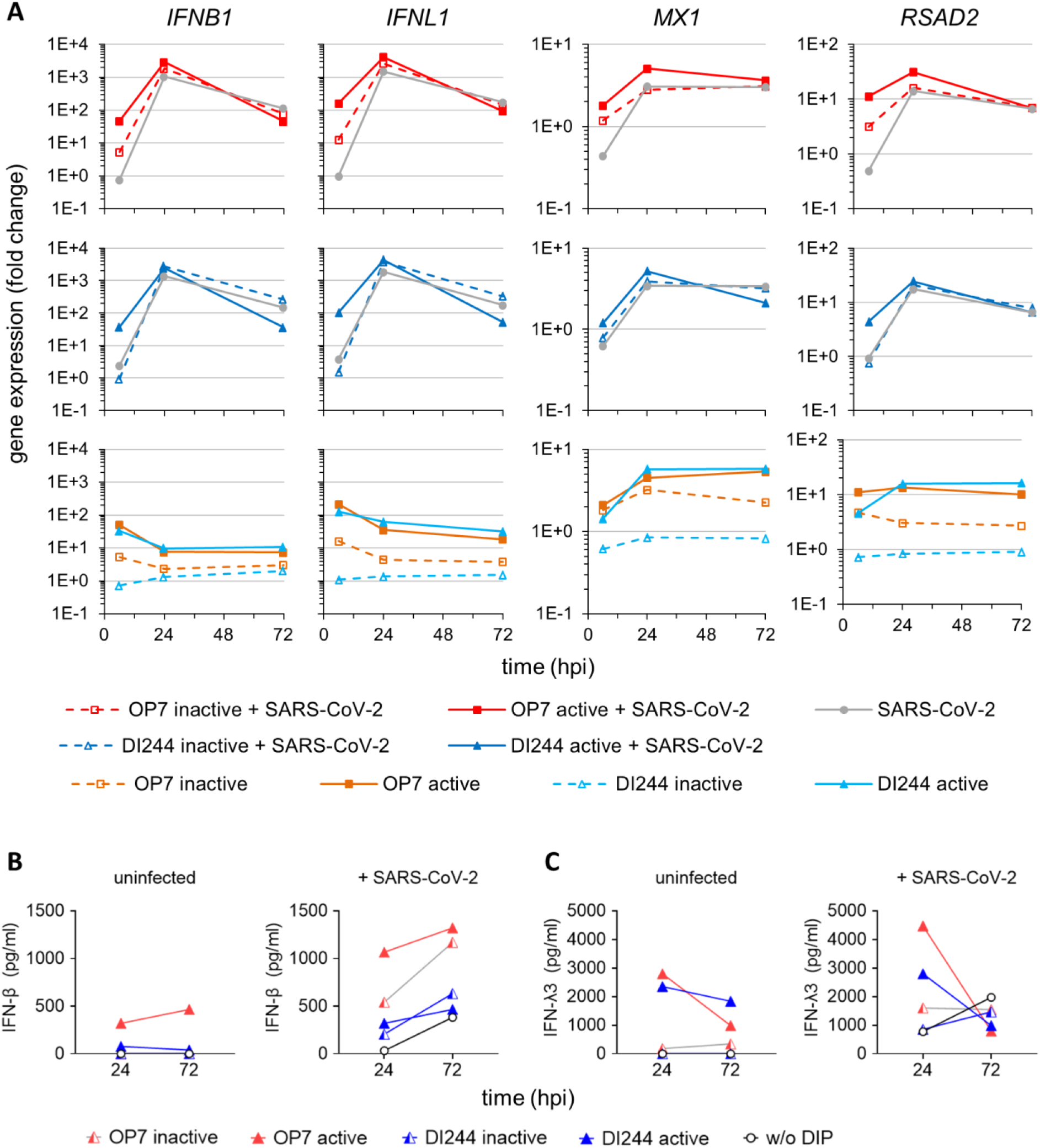
Stimulation of IFN-induced antiviral activity by IAV DIP infection. SARS-CoV-2-infected Calu-3 cells (MOI=0.03) were treated with IAV DIPs (DI244 or OP7) at 1 hpi. For DI244 and OP7 infection, 10% (v/v) (100 μL culture volume) of highly concentrated cell culture-derived DIP material was used (Hein et al., 2021c, Hein et al., 2021a). At indicated times post infection, infected cells were lysed to allow for total RNA extraction, required for (**A**,) gene expression analysis. In addition, supernatants were sampled for (**B** and **C**) quantification of secreted IFNs. Illustration includes data from one experiment. (**A**) Gene expression analysis of SARS-CoV-2 and IAV DIP co-infection. Transcript levels were quantified by real-time RT-qPCR and expressed as fold change (relative to untreated, uninfected cells). (**B** and **C**) Host cell IFN production. Protein levels of IFN-β (**B**) and IFN-λ3 (**C**) were assessed using ELISA.

**Fig. 4.**
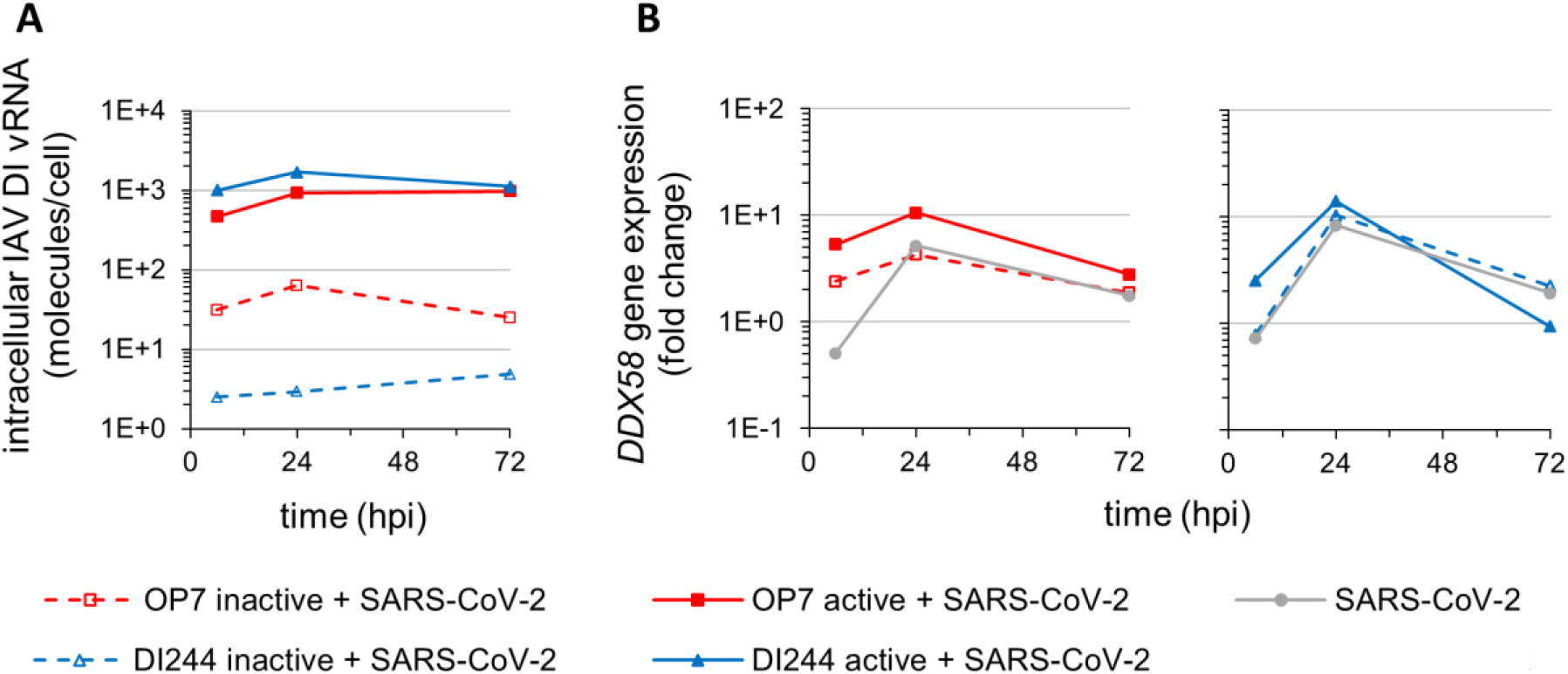
Cellular uptake of IAV DI vRNAs and *DDX58* expression. SARS-CoV-2-infected Calu-3 cells (MOI=0.03) were treated with IAV DIPs (DI244 or OP7) at 1 hpi. For DI244 and OP7 infection, 10% (v/v) (100 μL culture volume) of highly concentrated cell culture-derived DIP material was used (Hein et al., 2021c, Hein et al., 2021a). At indicated times post infection, cells were lysed to allow for total RNA extraction, required for (**A**) quantification of intracellular DI vRNAs and (**B**) analysis of *DDX58* gene expression. Illustration includes data from one experiment. (**A**) Intracellular DI vRNA levels during SARS-CoV-2 and IAV DIP co-infection. Cells were assayed for viral RNAs by real-time RT-qPCR. (**B**) *DDX58* gene expression analysis. *DDX58* (encoding for RIG-I) transcript levels were quantified by real-time RT-qPCR and expressed as fold change (relative to untreated, uninfected cells).

In line with the results from real-time RT-qPCR, protein levels of secreted IFN-β and IFN-λ3 (investigated using ELISA) were elevated at later times post infection for cells co-infected with IAV DIPs and SARS-CoV-2 compared to cells infected with only SARS-CoV-2 (Fig. 3B and 3C). In conclusion, our results suggest the early stimulation and subsequent boosting of the type-I and type-III IFN response as causative for the inhibiting potential of IAV DIPs against SARS-CoV-2 replication.

## Discussion

Despite the recent availability of vaccines against COVID-19, options for antiviral treatment are urgently needed for therapeutic application. Here, we show that cell culture-derived IAV DIPs are highly potent inhibitors of SARS-CoV-2 replication in human lung cells. In addition, our data obtained in *in vitro* experiments suggest that suppression of SARS-CoV-2 replication by IAV DIPs is predominantly attributed to their ability to stimulate innate immune responses ultimately inducing an antiviral state in target cells.

In the clinic, already approved antivirals for treatment of COVID-19 showed only very limited efficacy. For instance, treatment with the polymerase inhibitor remdesivir did not result in an overall decrease in mortality (Beigel et al., 2020, Pan et al., 2020). For patients receiving supplemental oxygen, however, an improvement in the survival rate from about 4% to 12% was observed (Beigel et al., 2020). In addition, the time required to recover from COVID-19 was decreased by five days (Wang et al., 2020, Beigel et al., 2020). Another option to treat COVID-19 is the use of monoclonal antibodies that target the receptor binding domain of the SARS-CoV-2 S protein, thereby inhibiting engagement with the host cell entry receptor angiotensin-converting enzyme 2 (ACE2) (Hoffmann et al., 2020, Abraham, 2020). Here, it was suggested to use antibody cocktails to prevent the emergence of viral escape variants in treated individuals (Baum et al., 2020). In clinical trials, treatment of outpatients with one such an antibody cocktail (i.e., bamlanivimab) accelerated the decrease in viral load and reduced the fraction of patients requiring hospitalization from 6.3% to 1.6% (Chen et al., 2020b). The administration of the corticosteroid dexamethasone (in clinical use) resulted in an overall lower mortality in critically ill COVID-19 patients (Horby et al., 2020, Sterne et al., 2020). This has a caveat, though, as a decrease in mortality was observed for patients requiring oxygen (including mechanical ventilation), but an increase in mortality was reported for patients not requiring oxygen (Horby et al., 2020). This comparatively little progress in COVID-19 therapy has sparked calls to invest more research into broadly acting antiviral agents against SARS-CoV-2 to combat future pandemics (Dolgin, 2021).

Treatment of COVID-19 patients with IFNs has not been approved yet. In general, SARS-CoV-2 infection can modulate and inhibit the IFN response (Chen et al., 2020a, Konno et al., 2020, Lei et al., 2020). In addition, it was recently shown that the host cell entry receptor ACE2 is indeed an ISG, and it was speculated that SARS-CoV-2 may exploit the IFN-driven upregulation of ACE2 to enhance infection (Ziegler et al., 2020). However, SARS-CoV-2 replication is, in general, also susceptible to inhibition by exogenously added IFN. For instance, all IFNs (type I, II and III) exhibited potent antiviral activity with SARS-CoV-2 replication *in vitro* (Busnadiego et al., 2020, Felgenhauer et al., 2020), suggesting that the antiviral activities of IFNs can counterbalance any proviral effects derived from ACE2 induction. Yet, note that type-I IFN treatment seems to play an ambiguous role in COVID-19, depending on the stage of the disease (reviewed in (Lee and Shin, 2020)), and also type-III IFNs can contribute to COVID-19 pathogenesis (Broggi et al., 2020). Therefore, therapeutic administration of IFNs (or IAV DIPs) in the future will have to be carefully evaluated with respect to timing. Nevertheless, it was shown that intranasal IFN-I administration (in hamsters) pre- or post-virus challenge can reduce SARS-CoV-2 disease burden (Hoagland et al., 2021). Moreover, in a placebo-controlled phase 2 clinical trial, administration of inhaled, nebulized IFN-β (to patients already admitted to hospital due to COVID-19 symptoms) resulted in a higher chance of disease improvement and a more rapid recovery from COVID-19 (Monk et al., 2020).

In our cell culture experiments, IAV DIPs completely abrogated SARS-CoV-2 replication. Notably, the UV-irradiated and thus inactive DIP material (containing degraded DI vRNAs) also showed a residual inhibitory effect. Yet, the observation of a much stronger antiviral effect upon treatment with active DIPs hints to an immunostimulatory activity of active IAV DIPs in the context of SARS-CoV-2 suppression. In principle, DIPs are defective in virus replication; therefore, they fail to complete the entire infection cycle. Thus, the genomic DI vRNAs entering the cell cannot multiply, but are typically very well detected by RIG-I (Baum and Garcia-Sastre, 2011), which subsequently leads to the activation of an IFN-response (Rehwinkel et al., 2010), in line with our results. In conclusion, it appears that physically (or chemically) inactivated viral particles are not sufficient to induce a strong antiviral immunity, in contrast to “live” but propagation-incompetent IAV DIPs. With respect to the safety of IAV DIPs for potential clinical application, it is important to note that administration of active DIPs did not cause apparent toxic effects in mice and ferrets (Hein et al., 2021a, Hein et al., 2021c, Dimmock et al., 2008, Dimmock et al., 2012).

Our results support the notion that IAV DIPs do not only protect host cells from IAV infection but, in addition, may generally confer protection against other heterologous IFN-sensitive respiratory viruses (Easton et al., 2011, Scott et al., 2011, Dimmock and Easton, 2015). Considering the emergence of new SARS-CoV-2 variants, against which a decreasing efficacy of various vaccines is becoming evident, the unspecific stimulation of innate immunity by IAV DIPs is considered advantageous; in particular, regarding a potential universal efficacy against such new (and future) variants. Furthermore, previous *in vitro* and *in vivo* experiments also revealed an antiviral effect of IAV DIPs against a variety of different IAV subtypes, including pandemic and highly pathogenic avian IAV strains (Dimmock et al., 2008, Dimmock et al., 2012, Zhao et al., 2018, Huo et al., 2020a).

Future work to clarify the therapeutic effects of IAV DIPs on the outcome of SARS-CoV-2 infection and to decipher in more detail the underlying mode of action should comprise animal trials in Syrian hamsters or genetically modified humanized K18-hACE2 mice, which are susceptible to SARS-CoV-2 infection and develop a similar respiratory disease compared to human COVID-19 (Kaptein et al., 2020, Boudewijns et al., 2020, Chan et al., 2020, Yinda et al., 2021). Animal experiments will help to elaborate on the potential applicability of IAV DIPs (e.g., administration via a droplet spray) as a pre- and post-exposure treatment for instance in acute SARS-CoV-2 outbreak scenarios in the clinics or geriatric institutions. In addition to vaccination, this would represent an interesting option for prophylactic treatment to boost antiviral immunity in persons at acute risk for an infection or for therapeutic treatment during an early phase post infection and as such may prevent fatal COVID-19 outcomes.

## Materials and methods

### Cells and viruses

Vero-6 cells (ATCC CRL-1586) were maintained in DMEM medium (Gibco, 4.5 g/L glucose, w/o pyruvate) supplemented with 10% fetal calf serum (FCS, Biowest, S1810-6500), 100 IU/mL penicillin, 100 μg/mL streptomycin, 1× GlutaMax (Gibco) and 1× sodium pyruvate (Gibco). Calu-3 cells (ATCC HTB-55) were cultured in MEM (Sigma) supplemented with 10% FCS (Biowest, S1810-6500), 100 IU/mL penicillin, 100 μg/mL streptomycin, 1× GlutaMax (Gibco) and 1× sodium pyruvate (Gibco). Caco-2 cells (ATCC HTB-37) were grown in MEM (Gibco) supplemented with 20% FCS (Biowest, S1810-6500), 100 IU/mL penicillin, 100 μg/mL streptomycin, 1× GlutaMax (Gibco) and 1× non-essential amino acid solution (Gibco). All cells were maintained or infected at 37°C in a 5% CO_2_ atmosphere.

The IAV DIPs DI244 and OP7 were produced in a cell culture-based process using a 500 mL laboratory scale stirred tank bioreactor, followed by purification and concentration by membrane-based steric exclusion chromatography (Marichal-Gallardo et al., 2017, Marichal-Gallardo et al., 2021), as described previously (Hein et al., 2021c, Hein et al., 2021a). Production titers of 3.3 and 3.67 log hemagglutination (HA) units/100μL (quantified by the HA assay (Kalbfuss et al., 2008)) and 5.6 × 10^8^ and 1.12 × 10^11^ DI vRNAs/mL (quantified by real-time RT-qPCR (Kupke et al., 2019, Hein et al., 2021c, Wasik et al., 2018)) were achieved for DI244 and OP7, respectively.

The SARS-CoV-2 isolate hCoV-19/Croatia/ZG-297-20/2020 was used. All experiments with infectious SARS-CoV-2 were performed in the BSL-3 facility at the Helmholtz Centre for Infection Research (Braunschweig, Germany). The SARS-CoV-2 seed virus was produced in Caco-2 cells, and virus particles were enriched in Vivaspin 20 columns (Sartorius Stedim, Biotech) via centrifugation. Collected virus was stored at −80°C. SARS-CoV-2 titers were quantified by plaque assay.

### SARS-CoV-2 quantification

Quantification of SARS-CoV-2 was performed by plaque assay. Samples were serially diluted in 10-fold steps, and used to infect a confluent monolayer of Vero-6 cells (on 96-well plates) for 1 h. Then, the inoculum was removed and cells were overlaid with cell culture medium containing 1.5% methyl-cellulose (SIGMA, #C9481-500). At 3 dpi, cells were fixed with 6% formaldehyde and stained with crystal violet. Wells were imaged using a Sartorius IncuCyte S3 (4× objective, whole-well scan) and plaque counts were determined.

### SARS-CoV-2 infection and antiviral treatment

Confluent Calu-3 cells in 96-well plates (~6 × 10^4^ cells/well) were infected with SARS-CoV-2 (2000 PFU per well). At 1 or 24 hpi, we added active or inactive IAV DIPs (DI244 or OP7) at indicated fractions (% v/v) with respect to the cell culture volume of 100 μL. Whenever indicated, we additionally added 0.8 μM ruxolitinib (Cayman Chemical, Cat. #Cay11609-1) to these wells. Alternatively, remdesivir (MedChem Express, #HY-104077) or human IFN-β-1A (PBL assay science, #11415-1) (instead of IAV DIPs) were added at indicated concentrations at 1 hpi. Supernatants were collected at indicated time points for quantification of SARS-CoV-2 titers (plaque assay) and for protein quantification of secreted IFNs using commercially available ELISA kits (see below). In addition, infected cells were lysed using solution “RL” for subsequent total RNA extraction using the “innuPREP RNA Mini Kit 2.0” (Analytik Jena, #845-KS-2040050), according to the manufacturer’s instructions, for gene expression analysis via real-time RT-qPCR.

### Immunofluorescence staining

(Co-)infected cells were fixed with 6% paraformaldehyde in PBS for 1 h at room temperature, followed by washing with PBS. Cells were permeabilized with 0.1% Triton X-100 in PBS for 10 min at room temperature, washed with PBS, and blocked with 2% BSA in PBS for 1 h. Antibody labelling was performed with mouse anti-SARS-CoV-2 S protein (Abcalis, clone AB68-A09, #ABK68-A09-M) and secondary antibody anti-mouse Alexa488 (Cell Signaling Technology, #4408), each step followed by three washing steps with PBS containing 0.05% Tween-20. Finally, cells were overlaid with Vectashield mouting medium (Biozol, #VEC-H-1000).

### Assessment of IFN production by host cells

Supernatants of (co-)infected cells were assessed for IFN-β and IFN-λ3 levels using corresponding “Quantikine” ELISA kits (R&D Systems, #DIFNB0 and #D28B00, respectively) according to the manufacturer’s instructions.

### Gene expression analysis

mRNA expression levels in (co-)infected cells were assessed using real-time RT-qPCR. 500 ng of total RNA was reverse transcribed using an oligo(dT) primer and the enzyme “Maxima H Minus” (both from Thermo Scientific) according to the manufacturer’s instructions. Next, qPCR was conducted using the “Rotor-Gene Q” real-time PCR cycler (Qiagen) and the following primers: *IFNB1*, 5’-CATTACCTGAAGGCCAAGGA-3’ and 5’-CAGCATCTGCTGGTTGAAGA-3’; *IFNL1*, 5’-GGTGACTTTGGTGCTAGGCT-3’ and 5’-TGAGTGACTCTTCCAAGGCG-3’; *MX1*, 5’-GTATCACAGAGCTGTTCTCCTG-3’ and 5’-CTCCCACTCCCTGAAATCTG-3’; *RSAD2*, 5’-CCCCAACCAGCGTCAACTAT-3’ and 5’-TGATCTTCTCCATACCAGCTTCC-3’; *GAPDH*, 5’-CTGGCGTCTTCACCACCATGG-3’ and 5’-CATCACGCCACAGTTTCCCGG-3’ (all sequences from (Busnadiego et al., 2020)); *DDX58*, 5’-TGCAAGCTGTGTGCTTCTCT-3’ and 5’-TCCTGAAAAACTTCTGGGGCT-3’ (Zhang et al., 2008). The qPCR reaction mixture (10 μL) comprised 1× Rotor-Gene SYBR green PCR mix (Qiagen), 500 nM of each primer, and 4 μL of cDNA. DNA denaturation was conducted for 5 min at 95°C, followed by 40 PCR cycles: 10 s at 95°C and 20 s at 62°C. Gene expression was calculated using the ΔΔCT method using *GAPDH* as the reference gene and expressed as fold change relative to untreated, uninfected cells.

### Quantification of intracellular IAV DI vRNAs

Real-time RT-qPCR was used for quantification of intracellular DI vRNAs from purified total RNA, as described previously (Hein et al., 2021a, Hein et al., 2021c, Kupke et al., 2019, Wasik et al., 2018). In brief, a primer system was used that allows polarity- and gene-specific detection of individual IAV vRNAs (Kawakami et al., 2011). To enable absolute quantification, RNA reference standards were synthesized and levels of vRNAs were calculated based on standard curves, as described previously (Kupke et al., 2019).

For RT, 1 μL of the total RNA sample was combined with 1 μL of dNTPs (10 mM) and 1 μL of the RT primer (1 μM) and filled up to a volume of 15 μL with nuclease-free water. Incubation was performed at 65°C for 5 min and 55°C for 5 min. During the latter step, a pre-warmed mixture (55°C) consisting of 4 μL of 5x RT buffer, 0.5 μL (100 U) Maxima H Minus reverse transcriptase, and 0.5 μL RiboLock RNase Inhibitor (all reagents from Thermo Scientific) was added. RT primer: OP7, 5’-ATTTAGGTGACACTATAGAAGCGACTGTGACTGCTGAAGTGGTG-3’; DI244, 5’-ATTTAGGTGACACTATAGAAGCGAGCGAAAGCAGGTCAATTATATTC-3’. RT was conducted for 30 min at 60°C, followed by 85°C for 5 min. Further, RNA reference standards in 10-fold dilution steps were reverse transcribed. Next, the cDNA reaction products were diluted to 100 μL in nuclease-free water. For qPCR, the Rotor-Gene Q real-time PCR cycler (Qiagen) was used. The qPCR mix (10 μL) contained 1× Rotor-Gene SYBR green PCR mix (Qiagen), 500 nM each primer, and 4 μL of diluted cDNA. qPCR Primers: OP7, 5’-ATTTAGGTGACACTATAGAAGCG-3’ and 5’-CATTTGCCTAGCCCGAATC-3’; DI244, 5’-ATTTAGGTGACACTATAGAAGCG-3’ and 5’-GGAATCCCCTCAGTCTTC-3’. The PCR cycling conditions comprised: initial denaturation step at 95°C for 5 min, followed by 40 PCR cycles of 95°C for 10 s and 62°C for 20 s.

## Supporting information

Full dataset (of Fig. 1-4)

## Acknowledgement

We thank Nancy Wynserski for excellent technical assistance. This research was supported by a grant from the German Federal Ministry of Science and Education (Grant No. 01KI20140A). P.M.G. kindly acknowledges joint funding from the German Federal Ministry for Economic Affairs and Energy (BMWi), the European Social Fund (ESF), and the Max Planck Society (MPG) within the *EXIST Forschungstranfer* program (project 03EFMST030).

## Author Contributions

Conceptualization, U.R. (Ulfert Rand), S.Y.K., L.C.S., U.R. (Udo Reichl) and D.B.; Investigation, U.R. (Ulfert Rand), H.S., M.D.H., T.H. and P.M.G.; Formal Analysis, U.R. (Ulfert Rand); Validation, U.R. (Ulfert Rand), H.S. and T.H.; Writing – Original Draft, S.Y.K.; Writing – Review & Editing, U.R. (Ulfert Rand), H.S., M.D.H., P.M.G., U.R. (Udo Reichl) and D.B.; Visualization, U.R. (Ulfert Rand) and S.Y.K.; Supervision, U.R. (Ulfert Rand), S.Y.K., L.C.S, U.R. (Udo Reichl) and D.B.; Project Administration, U.R. (Ulfert Rand), S.Y.K. and D.B.; Funding Acquisition, P.M.G., U.R. (Udo Reichl) and L.C.S.

## Data availability

Full dataset (of Fig. 1–4) is available in supplemental Table S1.

